# Genetic regulation of cell-type specific chromatin accessibility shapes the etiology of brain diseases

**DOI:** 10.1101/2023.03.02.530826

**Authors:** Biao Zeng, Jaroslav Bendl, Chengyu Deng, Donghoon Lee, Ruth Misir, Sarah M. Reach, Steven P. Kleopoulos, Pavan Auluck, Stefano Marenco, David A. Lewis, Vahram Haroutunian, Nadav Ahituv, John F. Fullard, Gabriel E. Hoffman, Panos Roussos

## Abstract

Nucleotide variants in cell type-specific gene regulatory elements in the human brain are major risk factors of human disease. We measured chromatin accessibility in sorted neurons and glia from 1,932 samples of human postmortem brain and identified 34,539 open chromatin regions with chromatin accessibility quantitative trait loci (caQTL). Only 10.4% of caQTL are shared between neurons and glia, supporting the cell type specificity of genetic regulation of the brain regulome. Incorporating allele specific chromatin accessibility improves statistical fine-mapping and refines molecular mechanisms underlying disease risk. Using massively parallel reporter assays in induced excitatory neurons, we screened 19,893 brain QTLs, identifying the functional impact of 476 regulatory variants. Combined, this comprehensive resource captures variation in the human brain regulome and provides novel insights into brain disease etiology.

**One sentence summary:** Cell-type specific chromatin accessibility QTL reveals regulatory mechanisms underlying brain diseases.

---

Genome-wide association studies (GWAS) have led to the identification of hundreds of genomic loci that are linked to increased risk for a range of brain diseases, including schizophrenia, Alzheimer’s disease, bipolar disorder and depression (*1*–*4*). However, due to linkage disequilibrium (LD), in which all genetic variants in the associated locus are strongly correlated, pinpointing causal variants to interpret GWAS hits is challenging (*5, 6*). In addition, the majority of identified common variants reside in non-coding regions of the genome and are enriched within regulatory elements, indicative of their impact on gene expression rather than on protein structure and function (*7*).

Gene expression quantitative trait loci (eQTL), have been used to facilitate interpretation of GWAS results (*8, 9*). However, eQTL-GWAS integration solely examines the relationship between genes and traits, and leaves a gap in understanding underlying regulatory mechanisms (*10*). Non-coding disease risk variants are thought to affect gene expression by modifying cell type-specific transcription factor binding to regulatory elements (*11, 12*). Since chromatin state is directly linked to transcription, the coexistence of similar chromatin accessibility quantitative trait loci (caQTL) and eQTL signals in a locus can reveal regulatory patterns linking risk variants to causal genes and disease mechanisms (*13*). Identifying disease-relevant regulatory elements can open up opportunities for targeted gene therapies with higher cell type specificity (*14*).

Efforts to better understand the precise etiology of neurodegenerative and neuropsychiatric diseases from a genetic perspective have been limited by the scale of cell-type specific data required. To date, only a few brain cell-type-specific caQTL studies have been conducted (*12*), including a recent study focusing on two cell-types, neurons and their progenitors, isolated from the dorsolateral prefrontal cortex (DLPFC) (*11*). Because of the limited statistical power of this particular study, only ∼2,000 caQTL were found across both cell types, accounting for just 2% of the open chromatin regions (OCRs). As such, further work is required to gain a more thorough understanding of the impact of regulatory mechanisms on gene expression and disease.

To investigate the genetic regulation of chromatin accessibility and its impact on neurodegenerative and neuropsychiatric diseases, we generated cell-type specific ATAC-seq data from neurons and glia isolated from 1,932 human postmortem brain samples derived from 616 donors and 4 distinct brain regions. Our analysis identified 34,539 OCRs with caQTL and revealed a high cell type specificity in the genetic regulation of chromatin accessibility. Integrating allele specific chromatin accessibility, eQTL and caQTL colocalization, and statistical fine-mapping, revealed putative molecular mechanisms for disease risk variants. We performed a massively parallel reporter assay (MPRA) in human induced pluripotent stem cell (hiPSC)-derived excitatory neurons, testing the functional impact of 19,893 candidate causal variants, and identified 476 variants with allelic effects. Overall, we provide a comprehensive catalog capturing variation in the human brain regulome, improving our understanding of the molecular basis of neurodegenerative and neuropsychiatric diseases.

## Results

### Cell-type specific regulation of chromatin accessibility

To characterize genetic regulatory variants underlying variation in chromatin accessibility in the human brain, we collected tissue samples from human postmortem brains. Tissue from the dorsolateral prefrontal cortex (DLPFC) and anterior cingulate cortex (ACC) were obtained from donors from the CommonMind Consortium (CMC), and samples from the superior temporal gyrus (STG) and parahippocampal gyrus (PHG) from the Accelerating Medicines Partnership Alzheimer’s Disease cohort (AMP-AD) (*15*) (**Fig. 1A**). Applying an established pipeline (*15, 16*), tissue samples were homogenized, nuclei were purified by fluorescence-activated nuclear sorting (FANS) using the neuronal marker NeuN, and chromatin accessibility was measured in neuronal (NeuN+) and glial (NeuN-) nuclei using ATAC-seq (*17*). Between the two cohorts, four brain regions, and two cell types, the dataset comprised 1,932 samples from 616 unique brain donors (**Fig. 1B**). Standard methods were applied for ATAC-seq processing, quality control and peak calling to identify OCRs in neurons and glia, covering 7.5% and 4.0% of the genome, respectively (*15*) (**Methods and Materials**).

**Figure 1.**
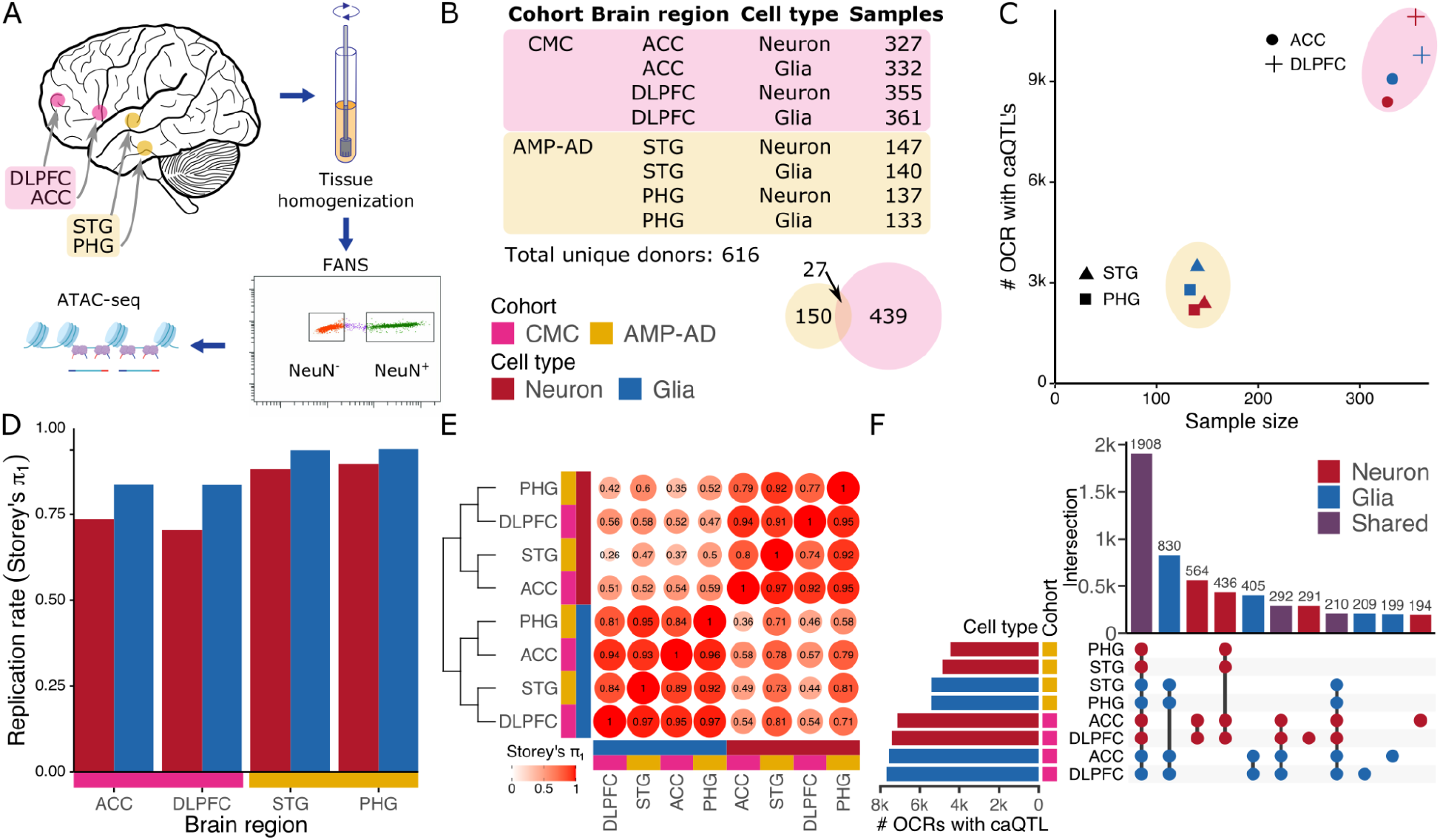
Cell-type specific regulation of chromatin accessibility. **A)** Experimental workflow including brain dissection, tissue homogenization, fluorescence-activated nuclear sorting (FANS) using the NeuN marker followed by ATAC-seq. **B)** Counts of donors and tissue samples for two cohorts, four brain regions and two cell types. Venn diagram shows donor count for each cohort along with a very small overlap. **C)** Number of OCRs with detected caQTLs compared to their sample size for each of 8 data subsets. Color indicates cell type, shape indicates brain region, and encompassing ellipse indicates cohort. **D)** Replication rate of caQTL’s discovered in each data subset in caQTL results from brain homogenate (*19*) using Storey’s π_1_ statistic (*22*). **E)** Replicate rate of caQTL’s across each pair of data subsets using Storey’s π_1_ statistic followed by hierarchical clustering. caQTL discovery was performed in the datasets along the columns and replication rate was measured in the datasets along the rows. **F)** Multivariate Bayesian meta-analysis identifies shared and cell type specific genetic regulatory effects on OCRs shared across cell types.

Using a linear mixed model to account for diverse genetic ancestry (*18*), genetic variants associated with chromatin accessibility within 50 kb of the OCR center were identified in 8 data subsets, stratified by brain region and cell type (*18*) (**Fig. 1C**). caQTL identified in this study showed high replication rates with those identified in data from brain homogenate (*19*) (**Fig. 1D**). Replication rates were higher for glia than neurons, consistent with their relative abundance in the cerebral cortex (*15, 20*). Evaluating the caQTL replication rates across 8 subsets of the data showed high replication rates for data subsets from the same cell type, and low replication rates across cell types (**Fig. 1E**). Analysis of cell type specific OCRs identified 14,529 OCRs in neurons and 15,860 in glia, with caQTL detected in at least one brain region. Analysis of 71,590 OCRs shared across cell types using multivariate Bayesian meta-analysis (*21*) identified 11,284 with a caQTL, including 2,033 (18.0%) with caQTL specific to neurons and 2,314 (20.5%) specific to glia (**Fig. 1F**).

### Cell type specific meta-analysis of caQTL signals

Since the difference between genetic regulation in neurons versus glia was the major axis of variation in the caQTL results, we performed fixed effect meta-analysis within each cell type, while statistically accounting for repeated measurements per donor (*18*). By further enhancing statistical power, analysis of cell type specific OCRs identified 11,475 of 303,992 (3.7%) OCRs in neurons and 8,890 of 197,553 (4.5%) in glia with genome-wide significant caQTLs (**Fig. 2A, S1**). Applying multivariate Bayesian meta-analysis (*21*) to the 96,138 shared OCRs detected in neurons and glia, identified only 3,603 (3.7%) of shared OCRs with caQTL detected in both cell types, while 4,761 (4.9%) and 5,810 (6.0%) had neuron- and glia-specific genetic regulation, respectively (**Fig. 2A**). Collectively, of the total 34,539 caQTLs detected in neurons and glia, only 10.4% are shared across cell types, strongly supporting the notion of cell type specific genetic regulation of brain OCRs.

**Figure 2.**
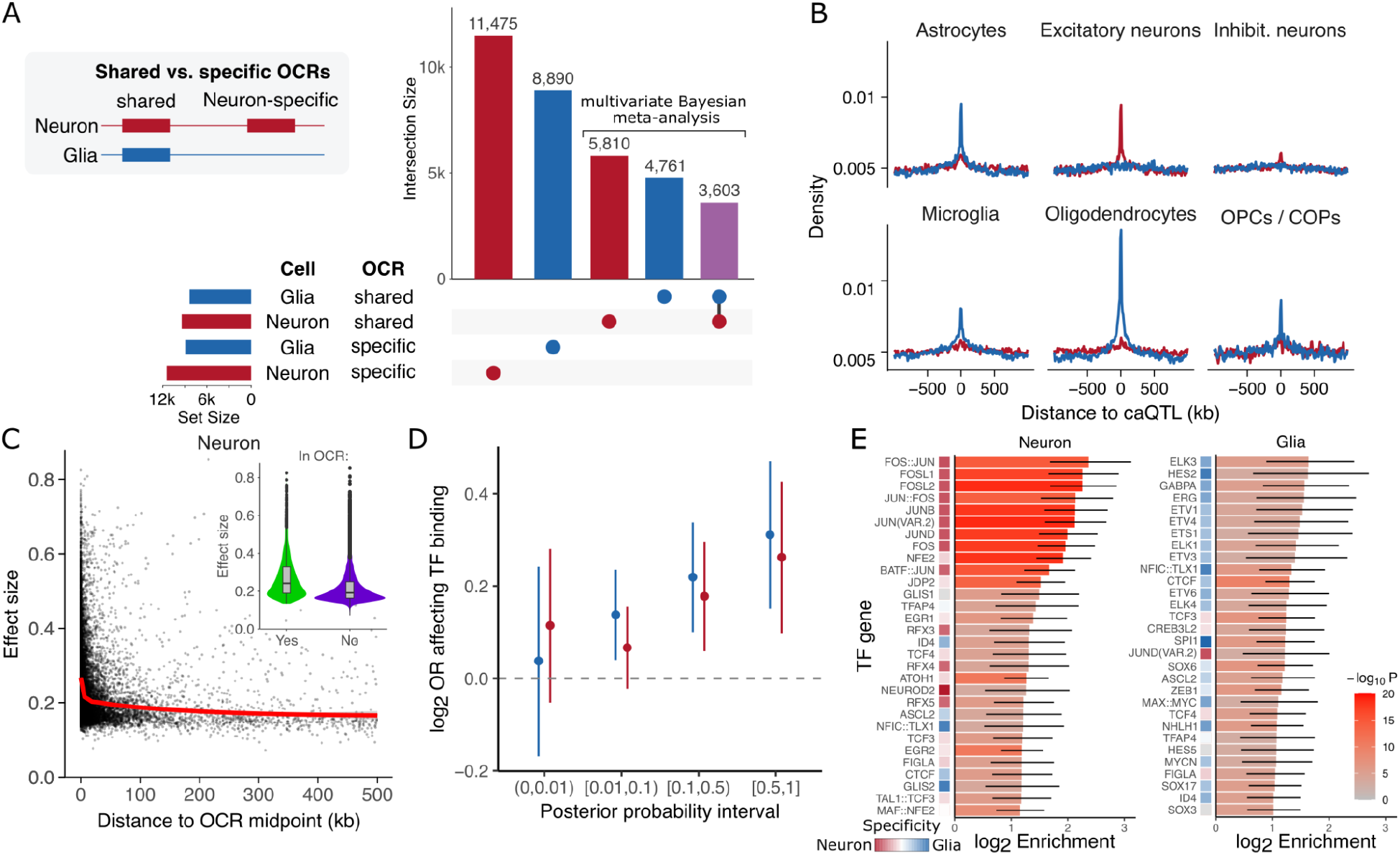
Characterizing cell type specific regulatory variants. **A)** Number of OCRs with detected caQTLs for cell type specific OCRs, and OCR shared across cell types. For the shared OCRs, the caQTL can have either a shared or cell type specific effect. **B)** Enrichment of lead caQTL variants within OCR discovered for 6 cell type populations from single cell ATAC-seq (*23*). **C)** Estimated effect size for detected caQTLs using an expanding search window up to 500 kb shows decay of effect size with distance. Inset shows larger effect size for variants within versus outside the OCR (p < 2.2e-16 by Wilcoxon test). **D)** Enrichment of fine-mapped variants within each posterior probability interval for disrupting a TFBS motif compared to the background set of tested SNPs not in the 95% credible set. **E)** Enrichment of fine-mapped variants in the 95% credible set for disrupting a TFBS motif shown for the top 30 transcription factors for each cell type. For each TF, its cell type specificity is shown based on the enrichment of its corresponding motif in neuronal- and glial-specific OCRs in the brain open chromatin atlas (*16*).

Evaluating enrichment of lead caQTL variants with OCRs identified from single nucleus ATAC-seq (*23*) supported the strong cell type specificity of the genetic regulatory programs (**Fig. 2B**). Glial caQTL were enriched in OCRs from astrocytes, microglia, oligodendrocytes and oligodendrocyte precursor cells, but not from neurons. Similarly, neuronal caQTLs showed the highest enrichment in excitatory neurons and were moderately enriched in inhibitory neurons, consistent with a higher abundance of excitatory neurons in the brain regions examined here (*15, 20*).

Genetic regulation of chromatin accessibility is mediated by variants modifying transcription factor binding sites (TFBS). Estimated caQTL effect sizes were significantly larger near the OCR where the majority of TFBS are present, and decay as the distance increases (**Fig. 2C, Fig. S2**). We performed statistical fine-mapping to identify candidate causal variants predicted to affect chromatin accessibility. We then integrated these results with variant-level computational predictions of TFBS disruption. Variants with higher posterior probabilities of affecting chromatin accessibility were enriched for predicted TFBS (**Fig. 2D**). Furthermore, motifs for specific transcription factors in neurons and glia showed higher enrichment for variants in the 95% credible set and acted in cell-specific manners (**Fig. 2E**).

### Allele specific chromatin accessibility

Heterozygous variants can provide additional information about genetic regulatory effects since an imbalance in the number of reads carrying the reference versus alternative allele indicates allele specific chromatin accessibility (ASCA) (**Fig. 3A**). The allelic ratio at each site was computed by summing all reads overlapping the variant across all samples for given cell type, and comparing to the null value of 50%. After applying strict quality control cutoffs and read filtering (**Methods and Materials**), 50.1% of reads corresponded to the alternative allele, indicating high quality ASCA analysis with minimal technical artifacts. In neurons, 13,816 of the 265,379 (5.2%) variants tested showed genome-wide significant ASCA and 21,937 of 183,159 (11.9%) were significant in glia (**Fig. 3B, S3**). Overlapping significant ASCA variants with OCRs from single cell ATAC-seq (*23*) identified a specific enrichment of neuronal variants in excitatory neuron OCRs, and enrichment of glial variants in oligodendrocyte and other non-neuronal OCRs (**Fig. 3C, S3**).

**Figure 3.**
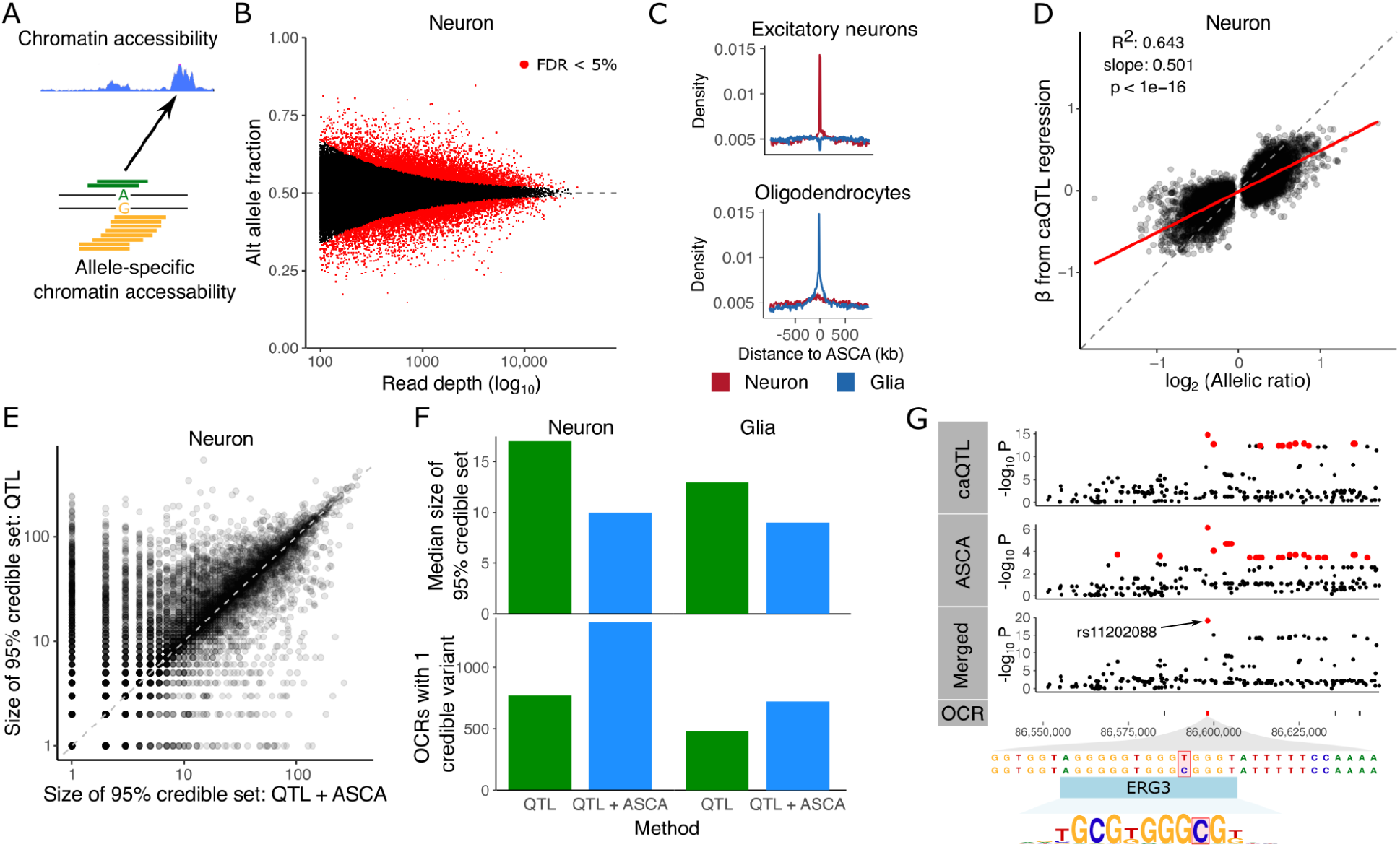
Allele specific chromatin accessibility. **A)** Diagram illustrating chromatin accessibility along the genome (top) and allele specific chromatin accessibility (ASCA) inferred by allelic imbalance. **B)** ASCA was inferred by testing the null hypothesis of equal fraction of alternative and reference alleles. Here, power to detect weak effects increases with the read depth. Red points indicate genome-wide FDR < 5%. **C)** Enrichment of significant ASCA variants in OCRs detected from single nucleus ATAC-seq (*23*). **D)** Relationship between the regression slope β estimated from caQTL regression and the allelic ratio from ASCA analysis. **E)** Comparison of the size of the 95% credible set from caQTL and merged (caQTL and ASCA) analyses. Each point represents an OCR. **F)** Comparison of fine-mapping results between caQTL and merged (caQTL and ASCA) analyses showing median size of credible sets (top) and number of OCRs with a single credible variant (bottom) for neurons and glia. **G)** Results for neuronal OCR using caQTL regression method, ASCA, and merged (caQTL and ASCA) analyses. Using the merged analysis reduces the 95% credible set to a single SNP, rs11202088, in the OCR that is predicted to disrupt an *ERG3* binding site. Variants in the 95% credible set are colored in red. OCR locations are shown and the target of the caQTL is colored red.

Standard regression-based caQTL analysis estimates the effect size as a slope, while ASCA estimates the effect based on allelic imbalance at heterozygous variants. The estimated effect sizes from caQTL and ASCA methods were highly correlated (R^2^ = 0.643, p<1e-16), consistent with both methods detecting genetic regulation effects using complementary aspects of the data (**Fig. 3D, S3**). The parameters of caQTL and ASCA testing are statistically independent under the null model of zero effect (*24*), which enables the combination of caQTL and ASCA results into a single test that increases statistical power while still controlling the false positive rate. We combined caQTL and ASCA signals (*25*) and then performed statistical fine-mapping on the combined results. Comparing the size of 95% credible sets from fine-mapping, combined caQTL and ASCA analysis yielded smaller credible sets for many OCRs compared to standard caQTL analysis, while showing high concordance between the two genome-wide (**Fig. 3E, S3**). The median size of the credible set reduced from 13 to 9 for glia and 17 to 10 for neurons, while the number of OCRs with a single credible variant increased from 481 to 723 in glia and 774 to 1,366 in neurons (**Fig. 3F**). For example, analysis of a neuronal OCR at chr10:86,597,844-86,598,604 using caQTL regression followed by fine-mapping nominates 14 SNPs in the 95% credible set, while ASCA nominates 25 SNPs. Merging these signals into a single statistical test leverages the advantages of both and substantially reduces the credible set to a single SNP, rs11202088, which is predicted to disrupt a ERG3 transcription factor binding motif (**Fig. 3G**).

### Genetic regulation is shared with gene expression

Genetic regulation of chromatin accessibility can drive downstream gene expression. We identified shared regulatory effects using colocalization analysis between caQTL and eQTL signals. Colocalization with eQTL results from brain homogenate (*18*) identified 2,569 gene-OCR pairs (involving 1,847 unique genes) for glia, and 2,012 gene-OCR pairs (involving 1,569 unique genes) for neurons with genetic regulation shared with chromatin accessibility in these cell types. Integrating with single nucleus RNA-seq (*26*) revealed strong cell type specificity of this shared genetic regulation. Neuronal caQTLs showed highest colocalization with eQTLs from excitatory neurons, while glial caQTLs showed higher sharing with oligodendrocytes, astrocytes, microglia and OPC/COPs (**Fig. 4A**).

**Figure 4.**
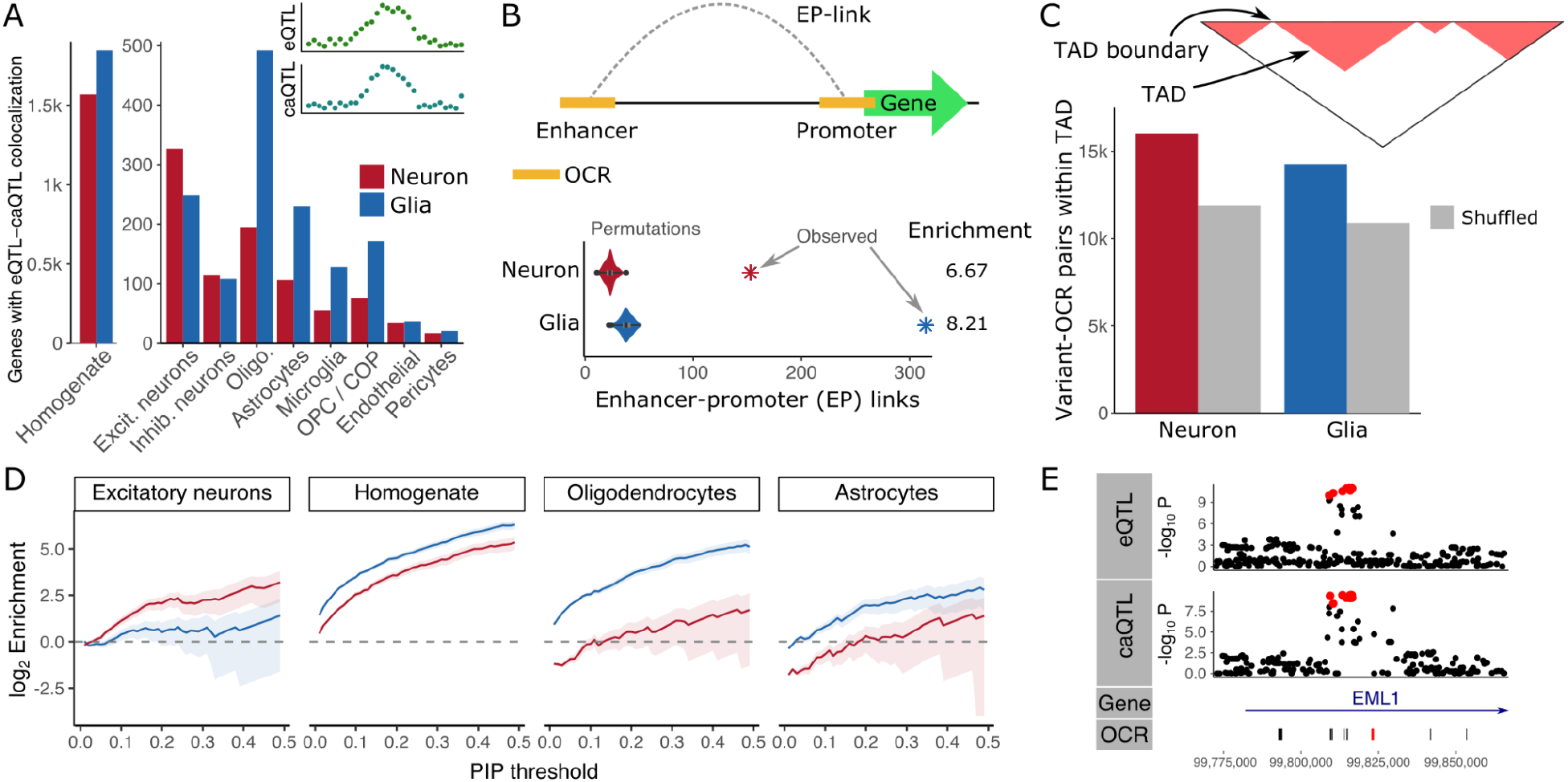
Shared genetic regulation of multiple molecular features. **A)** Number of genes in colocalized gene-OCR pairs for each cell type. **B)** OCR-gene pairs with shared genetic regulation are enriched for enhancer-promoter interaction. **C)** OCRs and their lead caQTL variant are enriched for being located in the same cell type specific topologically associated domain (TAD). **D)** Fine-mapped caQTL variants are enriched for fine-mapped eQTL variants from multiple cell types across a range of posterior probability thresholds. **E)** caQTL detected in glia for OCR at chr14:99,822,700-99,823,524 colocalizes with the eQTL signal for EML1 detected in brain homogenate (*18*). OCR locations are shown and the target of the caQTL is colored red.

Shared regulatory variants can act by increasing chromatin accessibility of a enhancer which then physically interacts with a gene’s promoter to drive expression (*27*). To test this mechanism in our data, we used the activity-by-contact method (*28*) to integrate our colocalized gene-OCR pairs with predicted enhancer-promoter links learned from cell type specific chromatin accessibility, histone modification and 3D genome profiling (*15*). Indeed, colocalized gene-OCR pairs were significantly enriched for having enhancer-promoter links in the corresponding cell type compared to the background of gene-OCR pairs with matched distance not sharing genetic regulation (p < 1e-16 for both cell types) (**Fig. 4B**). The structure of the 3D genome is also important for enabling genetic regulation of chromatin accessibility to have a downstream effect on expression. Using cell type specific 3D genome profiling, variant-OCR pairs were enriched for being in the same topologically associated domain compared to background (**Fig. 4C**).

Extending the colocalization analysis by integrating statistical fine-mapping on both chromatin accessibility and gene expression revealed strong cell type specificity of candidate causal variants (**Fig. 4D, S4**). Neuronal caQTL variants with high fine-mapping posterior probabilities were enriched for eQTL variants in excitatory neurons that also have high probabilities. Similarly fine-mapped glia caQTL variants were enriched in brain homogenate, oligodendrocytes and other cell types. For example, the caQTL signal in glia for an OCR at chr14:99,822,700-99,823,524 colocalizes with the genetic regulation of *EML1* identified in brain homogenate (*18*) (**Fig. 4E**).

### Integration of caQTL signal with disease risk variants

We further integrated fine-mapped caQTL variants from neurons and glia with risk variants for neuropsychiatric and neurodegenerative diseases, and other traits. Partitioning heritability for each trait using LD-score regression and estimating the contribution of many baseline variant annotations (*29*), variants in the 95% credible set for neurons and glia shows significant enrichments across a range of traits (**Fig. 5A**). Fine-mapped variants in glia, but not neurons, showed the strongest enrichment for Alzheimer’s disease risk variants, consistent with the role of microglia in the disease (*12, 30*). Enrichments are observed for both neurons and glia for schizophrenia and major depressive disorder (MDD), which is consistent with previous work (*23, 31*). Similarly, enrichments for both cell types are observed for neuroticism and education years since these traits have high co-heritability with schizophrenia (*32*).

**Figure 5.**
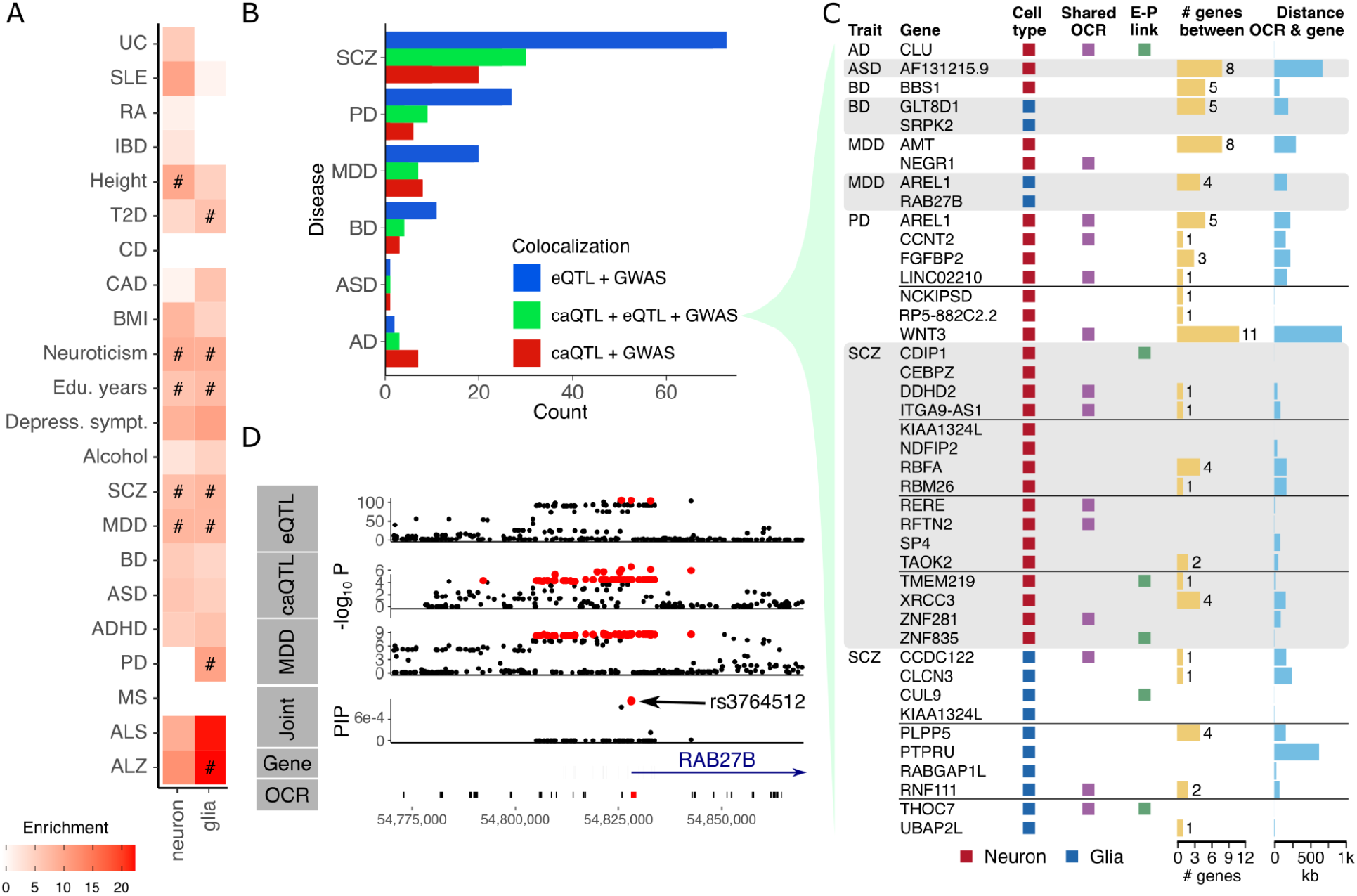
Integration of caQTLs with disease risk. **A)** Enrichment of caQTL variants in the 95% credible set for risk variants computed by LD-score regression (*29*). **B)** Counts of colocalized regions based on signals from caQTL, eQTL and GWAS for neuropsychiatric and neurodegenerative disease. Counts are stratified based on which signals colocalize. **C)** Colocalization between caQTL, eQTL and GWAS signals where the caQTL signal is cell type specific and posterior probability from MOLOC is > 0.9. Colors indicate cell type, whether the OCR is shared between cell types, and whether an enhancer-promoter (E-P) link is identified. For each gene-OCR pair, the distance between them and the number of intervening genes is shown. **D)** Colocalization of signals from eQTL, caQTL from neurons and risk for major depressive disorder (MDD) supports shared genetic regulation. Variants in the 95% credible set are shown in red. Joint statistical fine-mapping across three traits identified the SNP rs3764512 as the top candidate. This SNP is located within an OCR at the transcription start site of RAB27B.

Applying colocalization analysis to identify shared genetic regulation of chromatin accessibility, gene expression and risk for neuropsychiatric and neurodegenerative diseases, gave insight into the molecular mechanisms of disease. Across 6 diseases, we identified 98 colocalized caQTL-eQTL-GWAS triplets, and 53 colocalized caQTL-GWAS pairs without a shared eQTL signal (**Fig. 5B**). The majority of colocalized triplets, as well as eQTL-GWAS pairs, are observed for schizophrenia due to the large sample size and large number of risk loci (*1*). Examining the triplets involving cell type specific caQTL effects with the strongest colocalization score reveals known risk loci, but identifies specific genes, OCRs and cell types driving disease risk (**Fig. 5C, Table S1**). For example, a cell type specific triplet in neurons implicates *RAB27B*, a GTPase involved in synaptic transmission (*33*), in the molecular etiology of MDD. Joint statistical fine-mapping between three traits nominated the SNP rs3764512 as the top candidate, which is located in a neuronal OCR at the gene’s transcription start site (**Fig. 5D**).

### High throughput functional characterization of regulatory variants

While identifying candidate causal variants from statistical fine-mapping is an important step toward understanding molecular mechanisms of regulatory genetics, experimental validation of these computational predictions is needed. We performed a massively parallel reporter assay (MPRA) in hiPSC-derived excitatory neurons (*34*) to test the allelic effects of candidate causal variants identified from our previous large-scale eQTL analysis of human postmortem brain homogenate (*18*) (**Fig. 6A**). We selected 19,893 SNPs with posterior probability > 1% for testing allelic effects. We also selected 2,000 SNPs in an OCR but with no eQTL signal, and 2,000 SNPs near a strong candidate but not in the credible set as negative controls. Of the selected SNPs 2,165 were also fine-mapped caQTL in neurons with posterior probability > 1%.

**Figure 6.**
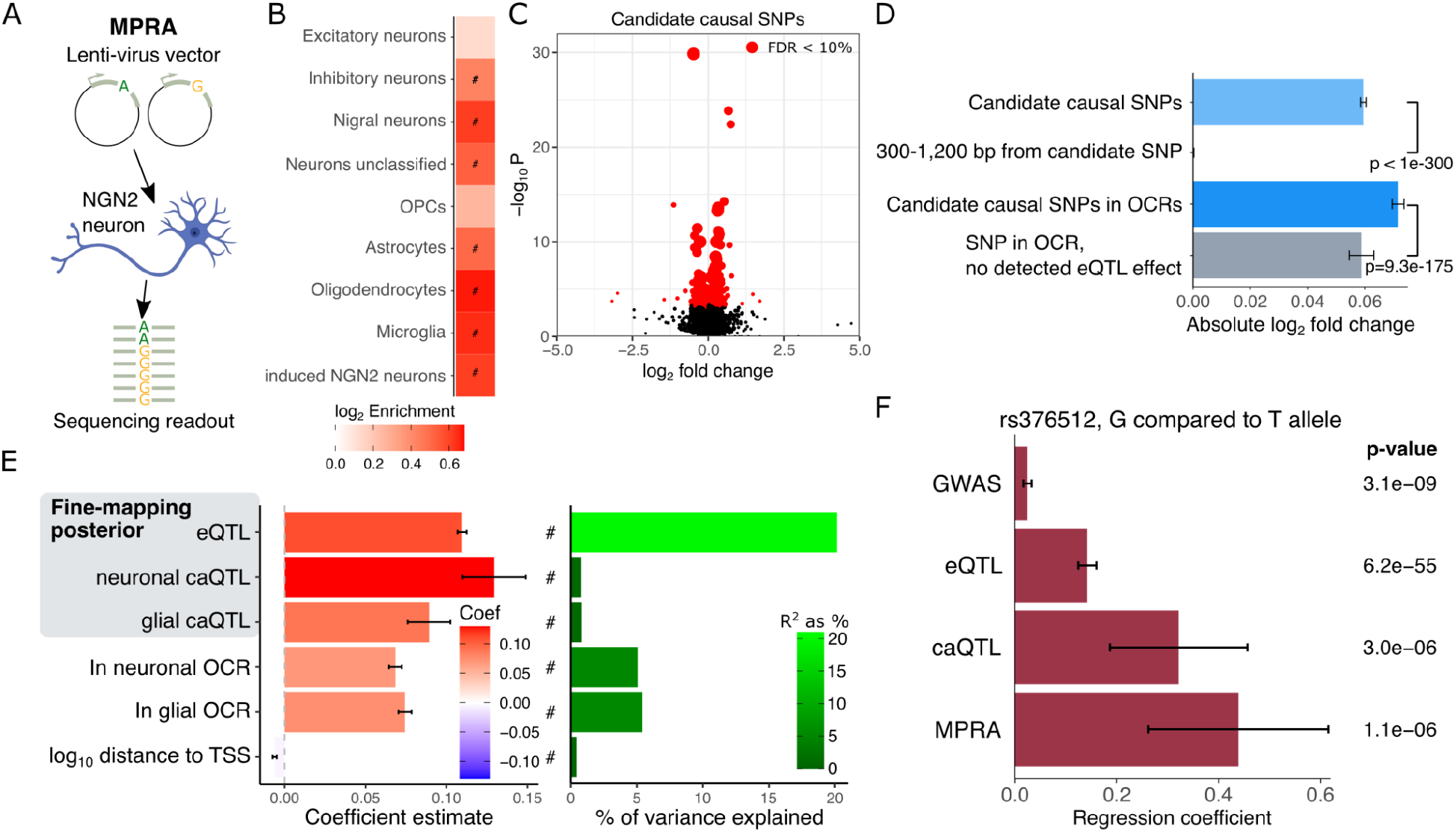
Massively parallel reporter assay. **A)** Diagram of massively parallel reporter assay (MPRA) with synthesized DNA constructs with two alleles inserted into a lentivirus MPRA vector. Vectors were transduced into human induced pluripotent stem cell (hiPSC)-derived excitatory NGN2 neurons, and the sequencing readout of DNA and RNA of barcodes was used to estimate differential allelic effects. **B)** Constructs with significant enhancer effects are enriched in open chromatin regions in multiple neuronal subtypes, in addition to non-neuronal cell types (*23, 36*). Enrichments with FDR < 5% are indicated with ‘#’. **C)** Volcano plot of allelic fold changes for candidates causal SNPs. SNPs with FDR < 10% are indicated in red, and larger points indicate SNPs with higher precision (i.e. lower standard error). **D)** Magnitude of allelic effects are shown for four sets of SNPs. Candidate causal SNPs have larger allelic effect magnitude than a SNPS 300-1,200 bp way. Candidate causal SNPs in OCRs have larger allelic effects than SNP in OCRs but with no detected eQTL. **E)** Regression coefficients showing relationship between magnitude of allelic effect and multiple SNP annotations (left). Color indicates the coefficient estimate. Associations with FDR < 5% are indicated by ‘#’. The percent of variation in allelic effect magnitude explained by each annotation is shown from the same regression models (right). **F)** Effect size estimates for rs3764512 comparing the G allele to the T allele shows consistency in direction.

Candidate regulatory sequences of 270 bp centered around each SNP were synthesized and cloned into a lentiMPRA construct (*34*). Reference and alternative alleles for each SNP were synthesized in separate sequences. The regulatory activity of each sequence was quantified by measuring the normalized RNA/DNA barcode abundance ratio (*34, 35*). High consistency was observed across three biological replicates (**Fig. S5**). To investigate the effects of sequence context, we additionally introduced SNPs at 25% and 75% positions of the construct for a subset of SNPs. Of the 994 variants tested at the three positions in the construct, 933 had sufficient sequencing converge in the MPRA. Of these, only 11 SNPs showed significant effect size heterogeneity at FDR < 5%, indicating there is little positional effect in our data (**Fig. S6**).

Testing each construct (i.e. 2 per SNP) for enhancer activity identified 1,267 sequences at FDR < 5%. These sequences were enriched in OCRs in multiple neuronal subtypes, non-neuronal cells and induced excitatory neurons (**Fig. 6B**). They were also enriched for known transcription factor binding sites motifs (**Fig. S7**). Testing constructs for allelic effects identified 476 SNPs at FDR < 10% (269 at FDR 5%) (**Fig. 6C**). The magnitude of the allelic effect is significantly higher for candidate causal SNPs than selected SNPs 300-1,200 bp away not in the credible set (**Fig. 6D**). Similarly, the effect magnitude for candidate causal SNPs within OCR is larger than SNPs in an OCR both with no eQTL signal.

Integrating the effect magnitude with SNP-level annotations reveals a strong relationship with the fine-mapping posterior probability from eQTL analysis (**Fig. 6E**). An increase in the posterior probability corresponded to an increase in effect magnitude. In fact, posterior probability explains 20.1% of the variation in effect magnitude, indicating that statistical fine-mapping is a good predictor of the functional impact of a SNP in this experiment. Posterior probabilities from caQTL in neurons and glia are also positively associated with effect magnitude, but explain much less variation since fewer tested SNP has non-zero posterior probabilities. Moreover, SNPs located within neuronal or glia OCRs had larger effect magnitudes that those outside, while SNPs further from corresponding gene’s transcription start site had a smaller effect magnitude.

Overall, we identified 7 variants with significant allelic effects from the MPRA that are associated with disease (**Table S2**). This small overlap is not unexpected since the MPRA was designed to test variants driving regulatory effects rather than disease risk. Nonetheless, intersecting the set of SNPs with experimentally validated allelic effects with colocalization and fine-mapping analysis from caQTL, eQTL and GWAS identified rs3764512, the top candidate for RAB27B in MDD. The G allele is the risk allele for MDD, and is predicted to drive an increase in chromatin accessibility, an increase in gene expression based on eQTL and increased enhancer activity from MPRA analyses (**Fig. 6F**).

## Discussion

Chromatin accessibility plays a pivotal role in the spatiotemporal regulation of gene expression. Previous studies indicate the importance of caQTL in the human brain (*19*) and available data shows that epigenetic regulation varies widely between different cell types, including in neurons (*11*) and microglia (*12*). Here, we present the largest study to date of the genetic regulation of chromatin accessibility in different cell types of the human brain. Combining signals across 4 brain regions increased statistical power, allowing us to identify 34,539 OCRs with genetic regulatory variants in both neurons and glia. Of these, only 10.4% were shared between them, a finding that is consistent with the diversity of open chromatin architecture across cell types in the brain (15, 16, 20, 23, 37).

Fine-mapping is a necessary step for understanding disease mechanisms and confidently identifying which downstream genes and pathways are affected (*6*). Yet solely using computational predictions from GWAS and statistical fine-mapping to overcome LD structure and identify context-specific causal variants remains challenging. In this study, we designed a step-wise approach to improve fine-mapping by combining direct analysis of neurons and glia isolated from human post mortem brain tissue with high throughput functional validation. Importantly, the discovery of cell type specific caQTL helps elucidate the molecular mechanisms driving disease risk. For example, Alzheimer’s disease shows a strong enrichment for fine-mapped variants in glia, while other neuropsychiatric diseases and traits (e.g. SCZ and MDD) show shared enrichment in both neurons and glia. Colocalization analysis identified cell type specific caQTL-eQTL-GWAS triplets with shared genetic regulation. These results nominate known and novel genes, OCRs and cell types driving disease risk.

We used a massively parallel reporter assay in hiPSC-derived excitatory NGN2 neurons to perform experimental validation of candidate causal variants identified in our large-scale eQTL analysis. We identified 476 variants with significant allelic effects, which is consistent with the number of variants validated with MPRAs in other studies (*38*–*40*). Overall, utilization of QTL fine-mapping served as a good predictor of the magnitude of allelic effects in the experimental model. For example, integrating MPRA results with colocalization analysis and fine-mapping implicated RAB27B, and the SNP rs3764512 in an OCR at its transcription start site, in the risk for MDD. RAB27B regulates exocytosis of dense-core vesicles in neuronal lines and synaptic vesicle release in neurons (*41*), which is relevant to the pathophysiology of MDD.

In conclusion, this study provides a population-scale human brain regulome resource capturing novel insights into brain disease etiology. Application of recent single cell technologies in future population-scale studies will increase resolution to further define cell type specific effects.

## Acknowledgements

We thank the patients and families who donated material for these studies. Brain tissue for the study was obtained through the NIH Neurobiobank from the following brain bank collections: The Mount Sinai/JJ Peters VA Medical Center NIH Brain and Tissue Repository, Brain Tissue Donation Program at the University of Pittsburgh and the Human Brain Collection Core (HBCC) within the National Institute of Mental Health’s Intramural Research Program (NIMH-IRP). We thank the computational resources and staff expertise provided by the Scientific Computing at the Icahn School of Medicine at Mount Sinai. We thank Dr. Pengfei Dong and other members in the Roussos Lab for helpful discussion.

## Author contributions

P.R. conceived of and designed the project. B.Z. led the caQTL analysis. J.B. led the processing and analysis of ATAC-seq data. C.D. performed the MPRA. D.L. and G.E.H designed the MPRA constructs. R.M., S.M.R. and S.P.K performed experimental work on brain tissue and data generation, supervised by J.F.F. Brain tissue specimens were provided by P.A., S.M., D.A.L., and V.H. N.A. supervised the MPRA. G.E.H. and P.R. supervised analysis. B.Z., J.B., J.F.F., G.E.H. and P.R. wrote the manuscript with input from all authors.

## Data and materials availability

The source data described in this manuscript are available via the PsychENCODE Knowledge Portal (https://psychencode.synapse.org/). The PsychENCODE Knowledge Portal is a platform for accessing data, analyses, and tools generated through grants funded by the National Institute of Mental Health (NIMH) PsychENCODE Consortium. Data is available for general research use according to the following requirements for data access and data attribution:

(https://psychencode.synapse.org/DataAccess). For access to content described in this manuscript see https://doi.org/10.7303/syn25955362.

