## Supplementary material for "Genetic regulation of cell-type specific chromatin accessibility shapes the etiology of brain diseases": Supplmental Methods and Materials

#### Supplementary Materials

Materials and Methods

Figs. S1 to S8

Tables S1 to S2

References

### Materials and Methods

#### Sample selection and preprocessing

We collected 1,949 ATAC-seq samples from two datasets generated previously in our lab at the Icahn School of Medicine at Mount Sinai (ISMMS). Both studies employed nearly identical experimental protocol and design, i.e. applying fluorescence-activated nuclear sorting (FANS) to isolate neuronal (NeuN+) and non-neuronal (NeuN-) nuclei from frozen brain tissue from two disease-relevant brain regions followed by ATAC-seq (*1*) to determine chromatin accessibility. From the study on Alzheimer's disease (AMP-AD dataset (*2*)), we obtained 557 samples from STG and PHG brain regions belonging to 177 donors (132 AD cases and 45 controls; 79 ATAC-seq samples were omitted due to the unavailability of genotype data). From the study on Schizophrenia and Bipolar disorder (CMC dataset (*3*)), we obtained 1,375 samples from ACC and DLPFC brain regions belonging to 466 donors (156 SCZ cases, 77 BD cases and 233 controls; 18 ATAC-seq samples were omitted due to the unavailability of genotype data).

All selected samples passed quality control criteria, including sex check and genotype check, as detailed in the original studies. They were processed by the same computational pipeline consisting of the following steps: (1) read trimming by Trimmomatic (*4*), (2) read mapping to human genome GRCh38 by the STAR aligner (*5*), (3) removing non-uniquely mapped reads and duplicated reads by PICARD (http://broadinstitute.github.io/picard/) and (4) peak calling by MACS2 (*6*). For the AMP-AD dataset, we prevented allelic bias resulting from person-specific genome variation by mapping on a custom reference genome (hg38) with masked biallelic SNPs detected from whole-genome sequencing (WGS) data of the AMP-AD donors. For the CMC dataset, we mapped on the standard version of the human genome (hg38 reference genome with the pseudoautosomal region masked on chromosome Y) using the WASP module (*7*) (available in the setting of STAR aligner) and supplied donors' WGS data. To perform peak calling in each dataset, we subsampled and merged all samples (BAM files) originating from the same cell type, brain region, and disease background. We then called peaks on those subsets and merged them into cell-specific consensus peaksets. After removing the peaks overlapping ENCODE blacklisted regions (*8*), we ended up with 315,630 neuronal OCRs and 205,120 glial OCRs in the AMP-AD samples, and 391,420 OCRs neuronal and 260,431 glial OCRs in the CMC samples.

#### Normalization on ATAC-seq peak signals

Before genetic association analysis, we performed normalization steps to remove hidden confounding factors. First, we implemented linear regression to regress out known biological and technical factors previously identified for the original datasets (*2*, *3*). While biological factors were chosen manually (i.e. Person ID, Brain region, Diagnosis status, Sex, Age), the technical factors were selected based on the Bayesian information criteria (BIC). Technical covariates varied across datasets as follows: (i) neuronal AMP-AD: GC_coverage_20-39%, (ii) glial AMP-AD: GC_coverage_20-39%, Fraction_of_reads_in_peaks, (iii) neuronal CMC:

GC_coverage_20-39%, GC_coverage_40-59%, GC_coverage_80-100%, Fraction_of_reads_in_peaks, STAR_unmapped_reads, AT_dropout, (iv) glial CMC: GC_coverage_0-19%, GC_coverage_20-39%, GC_coverage_80-100%, Fraction_of_reads_in_peaks.

To account for hidden covariates, we applied PEER (probabilistic estimation of expression residuals) (*9*) analysis. To find an optimal number of PEER factors, we performed multiple versions of caQTL analysis called on the input ATAC-seq matrix normalized by pre-selected biological and technical covariates but differing in the numbers of PEER factors between 4 and 28. Since the version of analysis utilizing 30 PEER factors produced the highest number of caQTLs (FDR<0.05) (**Fig. S8**), we used this number of factors to regress out the remaining hidden confounders from all ATAC-seq matrices.

#### Meta-analysis to integrate caQTL among data set

To improve the statistical power in QTL detection, we applied a meta-analysis method we have developed previously, mmQTL (*10*), and applied it on both eQTL and caQTL detection. mmQTL provides a flexible meta-analysis pipeline to integrate QTL among datasets, either individual-level or summary results. Briefly, it firstly performed QTL detection in each of the datasets, and assembled the QTL signal, of which non-significant variants were used to estimate covariance due to phenotypic correlation. This flexibility enables mmQTL to handle repeated measures from the same set of individuals while retaining high power and controlling the false positive rate. A linear mixed model was then used to estimate parameters.

#### Evaluating caQTL replicate rate

We used R package qvalue (*11*) to estimate caQTL replication rates. For a pair of datasets, we first extracted the most-significant variant for OCRs with caQTL in the discovery data. The p-values from the replication dataset were then used to estimate Storey's $\pi_1$ value, which indicates the fraction of hypothesis tests for which the null is rejected. Thus, $\pi_1$ is the estimated fraction of caQTLs that replicate in the second dataset. This metric of replication is useful because it does not depend on hard cutoffs for p-values for FDR, and it has been widely adopted.

#### Colocalization among eQTL, caQTL, and GWAS

To evaluate the relationship between molecular QTL, we used COLOC, an R package to conduct colocalization analysis (*12*). The output z-statistics from the meta-analysis was used as input for COLOC. The phenotypic variance was set to 1 as we have normalized the summary results before meta-analysis, but otherwise parameters were set to be default values. We also applied an extension, MOLOC (*13*), that applies this framework to identify colocalization of 3 signals. Here, we used it to identify shared genetic signals from meta-eQTL, meta-caQTL, and GWAS summary statistics. For COLOC analyses, we considered signals between two traits be colocalized at posterior probability ≥ 0.5 (i.e pp4≥0.5). For MOLOC considered signals between all three traits be colocalized at posterior probabilities ≥ 0.5 (i.e. pp14≥0.5).

#### Partitioned heritability

We applied a strategy developed in ref (*14*): for each meta-caQTL, we performed fine-mapping to compute the causal posterior probability (CPP) of each cis-SNP. Only those variants in the fine-mapped 95% credible set were retained for subsequent analysis. For each SNP in *cis*-regions, we assigned an annotation value based on the maximum value of CPP across all molecular phenotypes; SNPs that did not belong to any 95% credible set were assigned an annotation value of 0. This approach is referred to as MaxCPP in ref (*14*). StratifiedLD score regression (S-LDSC) was then used to partition trait heritability using the constructed functional annotations. The estimated enrichment was used to measure the importance of each caQTL category to human complex traits or diseases. To rule out the potential influences of the correlation among caQTL categories, we aggregated the baselineLD model, which includes a set of 75 functional annotations, to create functional annotations for the caQTL category and ran S-LDSR simultaneously.

We obtained GWAS summary statistics for 22 human complex traits or diseases, which contain both brain traits and non-brain traits. For brain traits, we included Alzheimer's disease (ALZ), Schizophrenia (SCZ), bipolar disorder (BIP), major depressive disorder (MDD), Parkinson's disease (PD) and multiple sclerosis (MS). We also downloaded summary results for non-brain traits or diseases, including height (HEIGHT), cardiovascular disease (CAD), Crohn's disease (CD), ulcerative colitis (UC) and rheumatoid arthritis (RA). The summary results were converted into the required format for LDSR by the provided "munge_sumstats.py" command in the LDSC package (https://github.com/bulik/ldsc).

#### Evaluation of the disruptive effect of caQTL on TF binding

We applied motifBreakR to annotate the disruptive effect of caQTL on TF binding (*15*). Briefly, fine-mapped caQTL variants located in ATAC-seq peaks among the 95% credible set were mapped to 630 known TF motifs from JASPAR2016 (*16*), and motifBreakR was used to calculate the relative entropy for the reference and alternative alleles. We only kept strongly affected TFBSs, and calculated the difference in each allele's binding ability.

To identify TFs more likely to be affected by caQTL, we randomly shuffled OCRs across the genome while retaining the number of OCRs and width distribution from the observed data. We then randomly selected variants for each OCR that matched the number and relative position of fine-mapped caQTL variants in the 95% credible set from the observed data. The number of variants affecting TF binding ability in the shuffled data was then computed to the observed data. Fisher's Exact test was utilized to calculate significance and FDR was controlled at 5% by the BH method. Shuffling was performed separately for each cell type to retain cell type specific properties of OCR and fine-mapped variants.

Cell type specificity of each TF was computed by evaluating the enrichment of the corresponding motif in neuron- or glia-specific OCRs from the Brain Open Chromatin Atlas (*17*), and scores were computed as described in Bendl, et al. (*2*).

#### Allele specific chromatin accessibility

From the sequence read mapping files, we used Genomic Analysis ToolKit (*18*) to calculate the allelic counts for each SNP within annotated OCRs, and to remove regions affected by potential mapping bias or other confounding factors. Each variant was required to have at least 2 reads mapped to each allele, and 3 heterozygous donors. A read depth of 100 was used to filter low confidence variants. Allele-specific chromatin accessibility was detected through the binomial test, and multiple tests were corrected by the p.adjust function in R, with FDR set to 5% via the BH method.

#### Allele-specific chromatin accessibility (ASCA) and caQTL

We applied the PLASMA strategy (*19*) to estimate ASCA. Imputed genotypes were first phased by SHAPEIT2 (*20*), and were used as input for Phaser (*21*) to estimate haplotypic expression from ATAC-seq BAM files. Haplotypic read counts were used by PLASMA to calculate ASCA, and the resulting summary results were then combined with meta caQTL signal, assuming independence between ASCA and caQTL. The sum of the test statistics from ASCA and caQTL was computed to a chi-square distribution with 2 degrees of freedom. These results were then used in downstream fine-mapping analysis.

#### Finemping on molecular QTL and GWAS

Statistical fine-mapping of caQTL and GWAS signals was performed with CAVIAR (*22*) using default parameters.

#### Multivariate Bayesian meta-analysis

We used MashR software (*23*) to assess the sharing of caQTL effects between different regions of the brain and cell types. MashR uses a Bayesian approach to estimate the posterior probability that each OCR in each cell type and brain region has a caQTL. By borrowing information across OCRs and conditions (i.e. cell types and brain regions) MashR can increase power to identify caQTLs, and also provide a direct test of caQTL sharing. The initial application of MashR considered 8 data subsets. Later analysis considered neuronal and glia signals which were previously combined across brain regions. Following MashR documentation, we used a set of ~90k OCRs shared by neuron and glia to estimate the prior effects size distribution, and ~600k randomly chosen variant-trait pairs were used to learn the empirical correlation structure.

#### Integration with other genomic annotations

Enhancer-promoter links were predicted with the activity-by-contact method (*24*) using our cell type specific ATAC-seq as well as cell type specific histone modification and 3D genome profiling as described by Bendl, et al. (*2*). The gene-OCR pairs with colocalized genetic signal were compared to gene-OCR pairs with matched distance not sharing genetic regulation. We generated 150 sets of matching gene-OCR pairs and used empirical null distribution to compute the enrichment and p-values.

Similarly, topologically associated domains (TADs) were identified from cell type specific 3D genome profiling as described by Bendl, et al. (*2*). The number of variant-OCR pairs within the same TAD was then evaluated, extending 500bp on each side to account for TAD-annotation uncertainty. Shuffled data was obtained by randomly relocating TAD start and end while retaining the width.

#### Design and analysis of MPRA

##### Sequences construct design

Variants tested in the MPRA were selected based on statistical fine-mapping from our previous eQTL analysis of 3,983 postmortem brain homogenate samples from 2,119 donors from multiple cohorts (*10*). The fine-mapping posterior inclusion probability (PIP) was used to select variants. A total of 39,786 constructs were used to test allelic effects. These were designed to be 270 bp long, and SNPs were placed in the center of the construct except for the exception described below.

**SNPs in candidate causal set (p1):**
The test set comprised 19,893 SNPs drawn from SNPs with PIP > 1% where the probability of selecting a SNP was proportional to its PIP. Using 2 alleles per SNP produced 39,786 sequences.

**Examining positional effects (p2):**
A set of 994 SNPs from set p1 was inserted 67 bp from the 3' end, and also 67 bp from the 5' end. Using 2 alleles in each of 2 positions produced 3,976 sequences.

**SNPs in an OCR, but without detected eQTL effect (n1):**
Set of 2,000 SNPs with no expected effect selected to have p-value > 0.1 to ensure no detected eQTL effect, minor allele frequency > 20% to ensure there was sufficient power to detect an effect, presence in an OCR detected in at least two cell types from single nucleus ATAC-seq of postmortem human brain (*25*). Using 2 alleles per SNP produces 4,000 sequences.

**SNPs near a strong candidate, but not in a credible set (n2):**

Set of 2,000 SNPs with no expected effect selected to have a detected eQTL signal at p<1e-8, within 300-1200bp of a SNP with PIP > 0.98, but not in the credible set. Using 2 alleles per SNP produces 4,000 sequences.

**Control sequences from previous literature (c1):**
We included previously characterized positive and negative sequences for NPC using lentiMPRA(*26*). We selected top 200 sequences with the highest enhancer scores and bottom 200 sequences with the lowest enhancer scores from fully differentiated hESC-derived NPC at 72 hour time point.

**Shuffled sequences for negative controls (c2):**
In addition, 100 shuffled sequences were included as negative controls for lentiMPRA.

#### Ngn2-inducible neuron culture

Human excitatory neurons were generated from an engineered hiPSC line WTC11 harboring a doxycycline-inducible neurogenin 2 (Ngn2) transgene at AAVS1 safe harbor locus. The differentiation was performed as previously described (*27*). Briefly, WTC11 that harbors an inducible Ngn2 expression cassette was dissociated with Accutase (07920, STEMCELL) following the Allen Institute hiPSC culture protocol, and seeded in Matrigel (354277, Corning)-coated plates using pre-differentiation medium supplemented with 10 µM ROCK inhibitor (S1049, VWR). The medium was prepared using KnockOut DMEM/F-12 (12660012, Fisher Scientific) with 1x N-2 (17502-048, Fisher Scientific), 1x NEAA (11140-050, Fisher Scientific), and freshly supplemented 10 ng/ml BDNF (450-02, PeProTech), 10ng/ml NT-3 (450-03, PeProTech), 1 µg/ml Laminin (23017015, Fisher Scientific), and 2 µg/ml freshly prepared doxycycline (D9891, Sigma). Medium without ROCK inhibitor was replaced daily for two days. Pre-differentiated neurons were then dissociated and subplated onto Poly-L-Ornithine-coated dishes with Maturation medium supplemented with 2 µg/ml doxycycline. Maturation medium was prepared by mixing Neurobasal-A (12349015, Fisher Scientific) with DMEM/F-12 (11330032, Fisher Scientific) at a 1:1 ratio with 1x NEAA, 0.5x B-27 (17504-044, Fisher Scientific), 0.5x N-2, 0.5x GlutMAX (35050061, Fisher Scientific) and freshly supplemented with 10 ng/ml BDNF, 10 ng/ml NT-3, and 1 µg/ml Laminin. This time point was counted as day 0. Half volume of medium was replaced without doxycycline at day 7 and 14 of differentiation.

#### LentiMPRA experiments

LentiMPRA was performed as previously described (*28*) with some modifications. Briefly, the designed oligo library was synthesized by Twist Bioscience. Two rounds of PCR were performed to add a minimal promoter and 15bp barcode downstream of the oligo. The PCR amplicon was cloned upstream of the GFP reporter gene in a lentiMPRA construct. The library was electroporated and amplified in electrocompetent cells (C3020K, NEB). Roughly 10 million colonies were harvested by midiprep (12945, Qiagen) to achieve an estimated 200 barcodes per oligo. Candidate sequences and barcodes were amplified from the plasmid library and sequenced by Illumina Nextseq mid-output PE150 to associate individual barcodes to candidate sequences. The Plasmid library was then packaged into a lentiviral vector using Lenti-Pac HIV Expression kit (LT002, GeneCopoeia) following the manufacturer's protocol. Crude viral solution was concentrated 100 times using Lenti-X Concentrator (631232, Takara Bio). Aliquots of lentivirus were immediately stored at -80°C. To determine lentiviral titer, $5 \times 10^4$ Ngn2 neurons were infected separately with 0,1,2,4,8,16,32 or 64 µl of lentivirus library at day 7. ViroMag R/L (RL41000, OZ Biosciences) was added to boost transduction efficiency. 6µl ViroMag per 100 µl lentivirus was used for optimized infection efficiency with minimal cell death. At day 14, DNA was extracted from infected cells and titer was calculated via qPCR. Based on the titration results, for each replicate, 7 million pre-differentiated neurons were plated in a 10cm dish and transduced with an estimated MOI of 90 at day 7 to reach an average of 65 random integrations per barcode. On day 14, DNA and RNA were harvested simultaneously using Allprep kit (80204, Qiagen), prepared into sequencing libraries via PCR and sequenced using Next high-output SE75.

#### Statistical analysis of MPRA

Barcode-candidate regulatory sequence (CRS) mapping and normalized barcode counting were performed using MPRAflow (*28*). The enhancer activity of CRS was compared against negative shuffled controls using R package MPRAnalyze (*29*). Testing of allelic effects was performed with MPRAnalyze applying their comparative framework. Benjamini-Hochberg multiple testing correction (*30*) was then applied.

Effect size heterogeneity was tested from the estimated logFC values and their standard errors using Cochran's Q-test implemented in metafor::rma() (*31*). Regression analysis with absolute log fold change as the response and fine-mapping posterior probability as a covariate was performed with precision weighting. Letting $se_i$ be the standard error of the $i^{th}$ log fold change, the weight for that entry is $1/ se_i^2$, following standing methods.

### Figs. S1 to S8

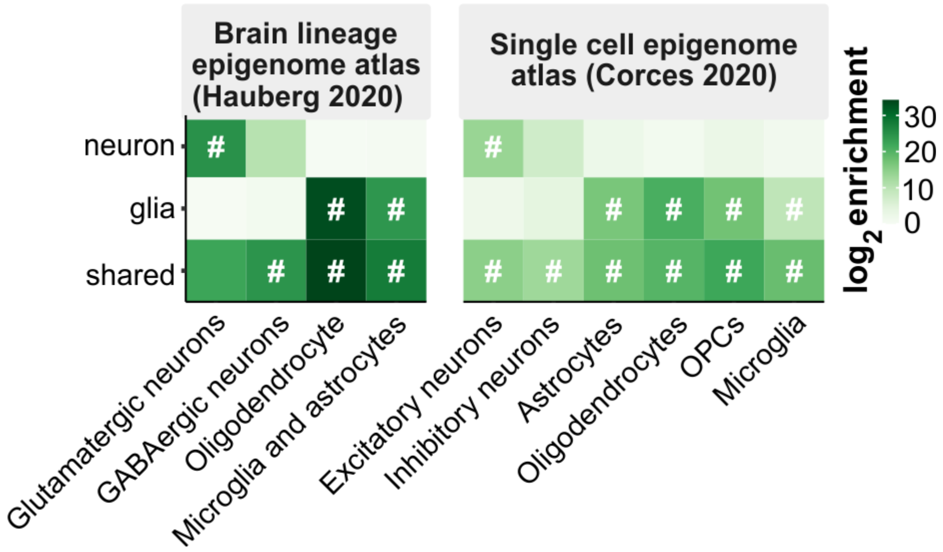


**Figure S1. Overlap of neuronal and glial OCRs with caQTL's with brain epigenome atlases** (*25*, *32*)**.** Enrichments with FDR < 5% are indicated with '#'.

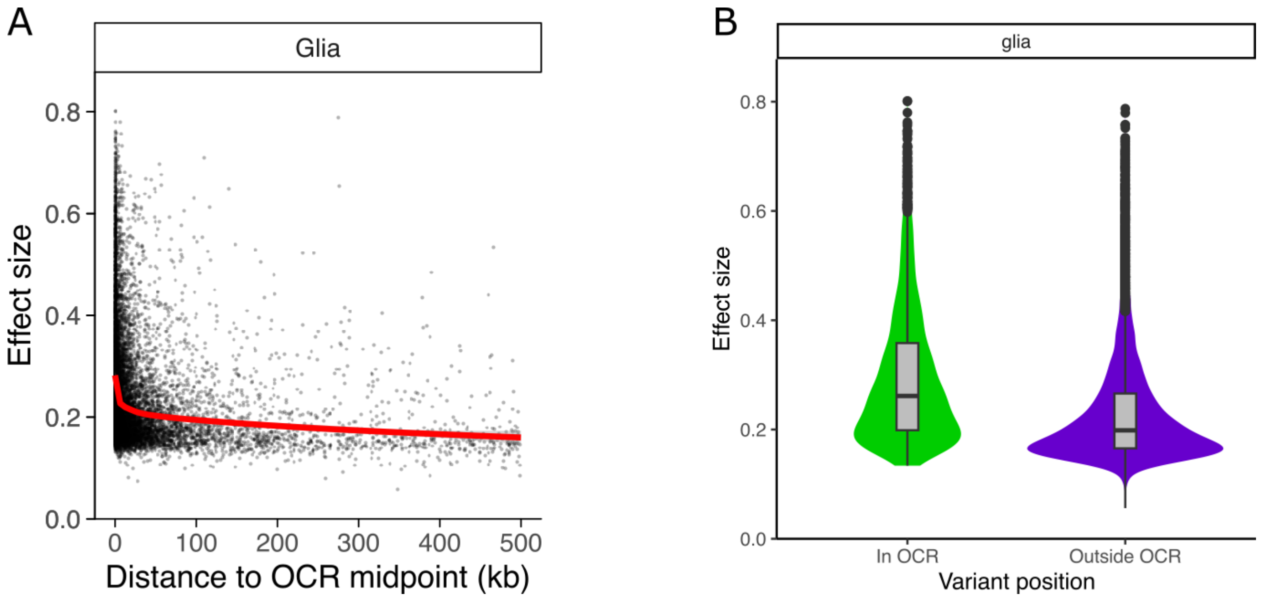


**Figure S2. Effect size of regulatory variants in glia.**
**A)** Estimated effect size for detected caQTLs using an expanding search window up to 500 kb shows decay of effect size with distance. **B)** Effect size is larger for variants within versus outside OCRs ($p < 2.2e-16$ by Wilcoxon test).

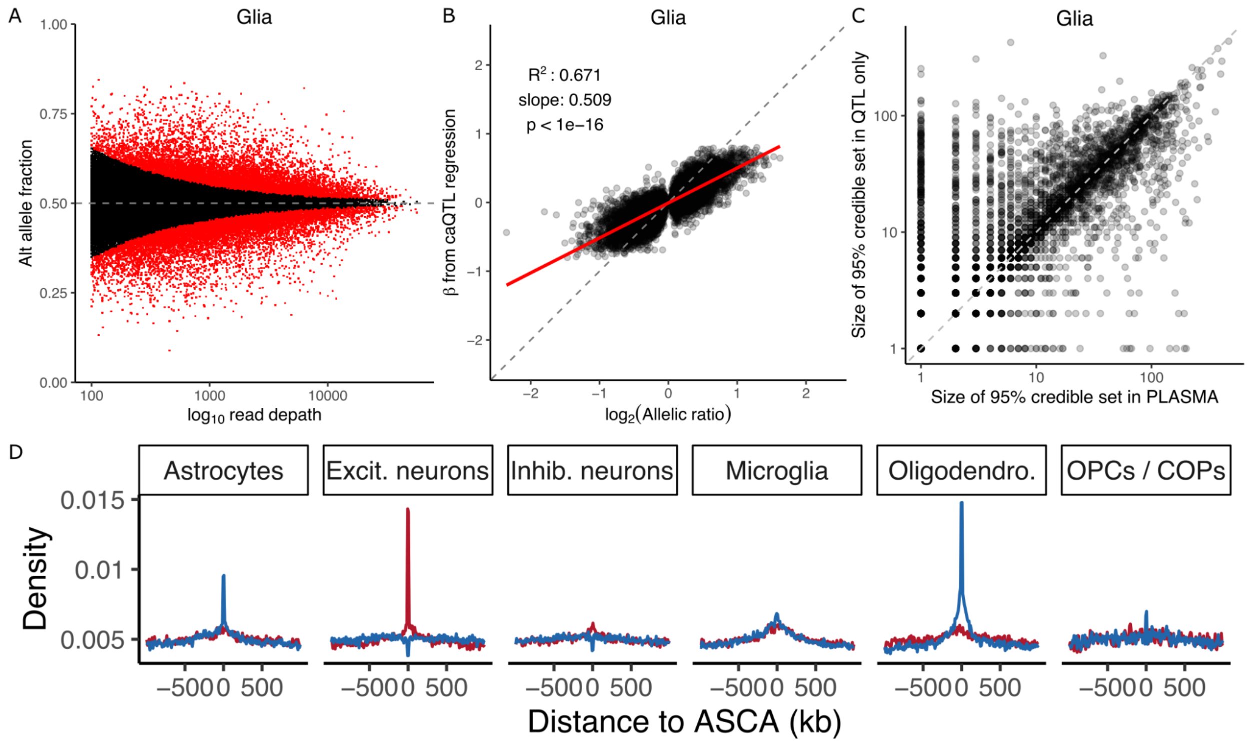


**Figure S3. Allele specific chromatin accessibility in glia.**

**A)** ASCA was inferred by testing the null hypothesis of equal fraction of alternative and reference alleles. Here, power to detect weak effects increases with the read depth. Red points indicate genome-wide FDR < 5%. **B)** Relationship between the regression slope β estimated from caQTL regression and the allelic ratio from ASCA analysis. **C)** Comparison of the size of the 95% credible set from caQTL analysis and caQTL + ASCA analysis. Each point represents an OCR. **D)** Enrichment of significant ASCA variants in OCRs detected from single nucleus ATAC-seq.

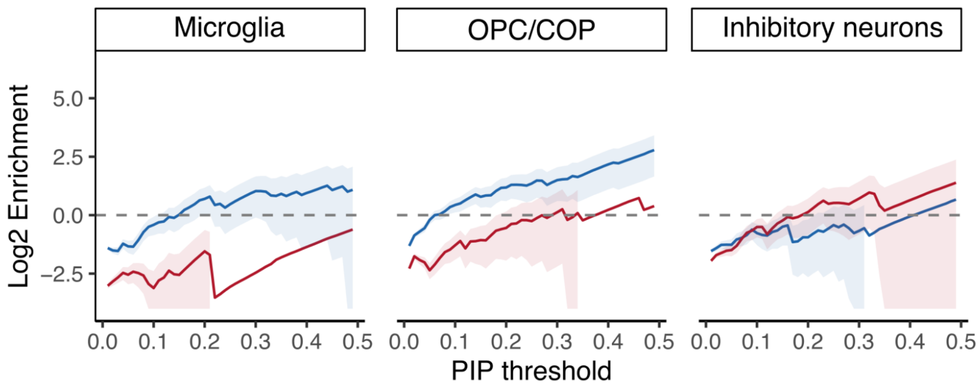


**Figure S4. Enrichment of fine-mapped variants in OCRs**

Fine-mapped caQTL variants are enriched for fine-mapped eQTL variants from multiple cell types across a range of posterior probability thresholds. Results shown here complement results in Figure 4D.

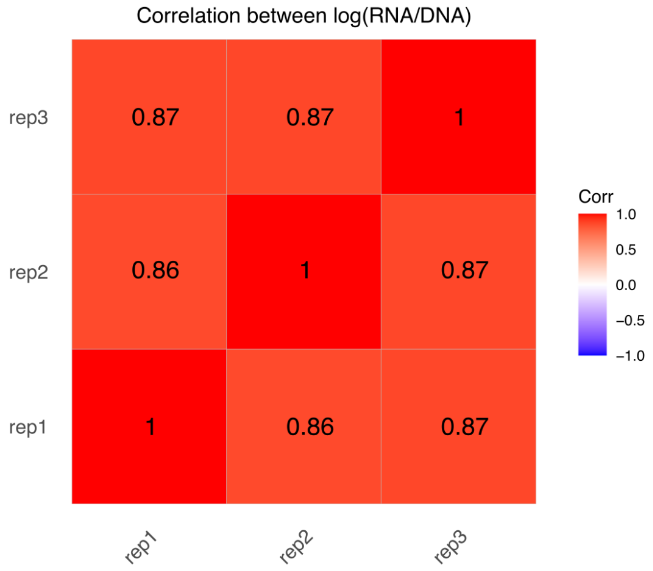


**Figure S5. Correlation between RNA and DNA log ratios from 3 technical replicates indicates high concordance.**

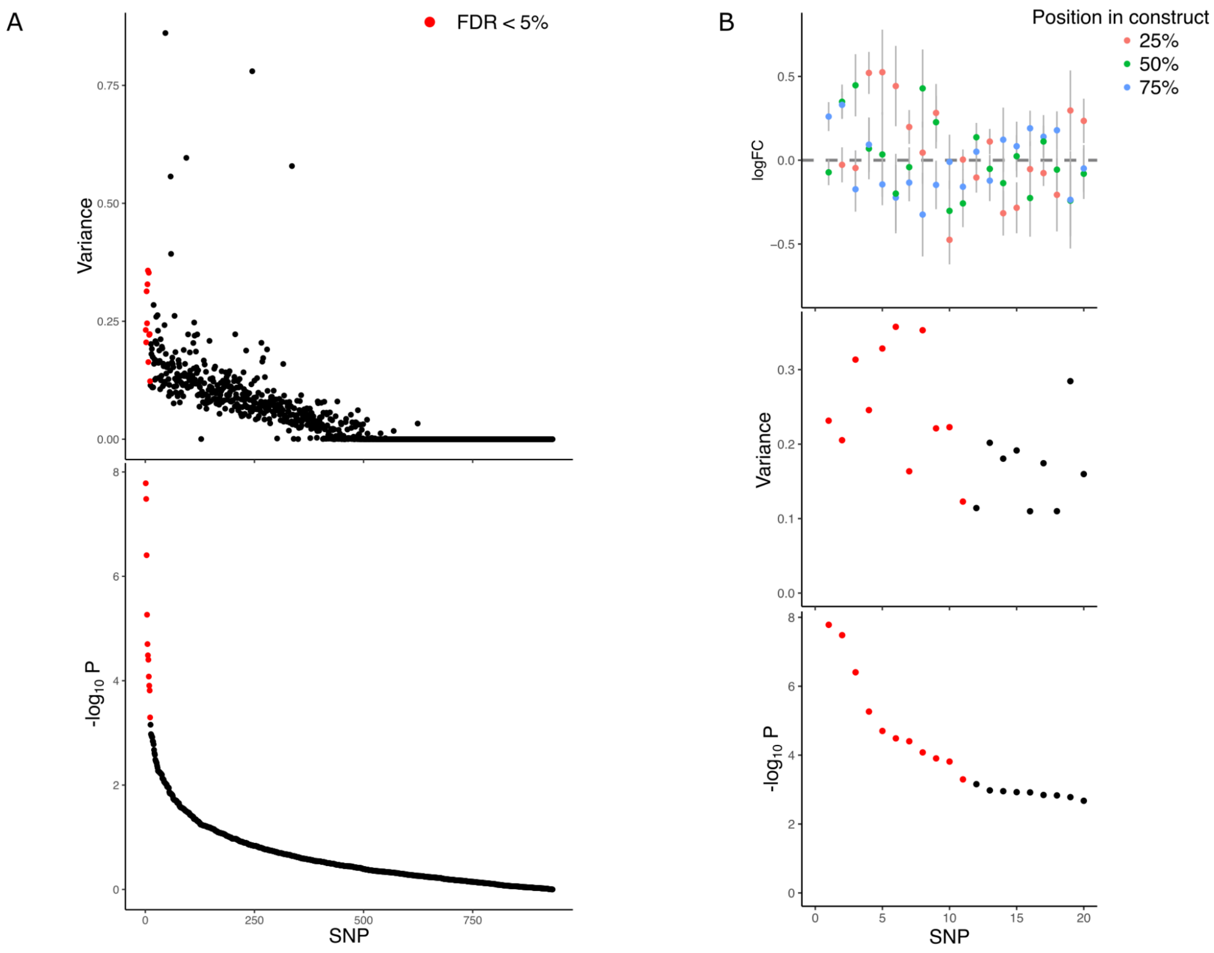


**Figure S6. Testing positional effects of variants by examining effect size heterogeneity.** Of the 994 variants tested for positional effect in the MPRA, 933 had observations for at least two positions that passed quality control. Of these 11 showed significant effect size heterogeneity across positions using Cochran's Q-test. **A)** Variance of the estimated effect size for each SNP (top) and their -$\log_{10}$ p-value from a test of effect size heterogeneity (bottom). **B)** For the 20 SNPs with the most significant effect size heterogeneity, we show the estimated effects and their 95% confidence interval (top). Colors indicate the position in the sequence relative to the 5' end. The variance in effect sizes (middle) and -$\log_{10}$ p-value from a test of effect size heterogeneity (bottom) are also shown.

| Motif | Name | p-value | q-value | # Target Sequences with Motif | % of Targets Sequences with Motif | # Background Sequences with Motif | % of Background Sequences with Motif |
|---|---|---|---|---|---|---|---|
| 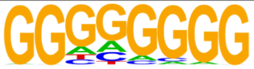 | Maz(Zf)/HepG2-Maz-ChIP-Seq(GSE31477)/Homer | 1e-10 | 0.0000 | 255.0 | 22.57% | 3070.8 | 15.12% |
| 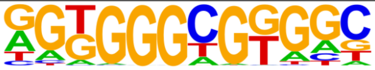 | KLF14(Zf)/HEK293-KLF14.GFP-ChIP-Seq(GSE58341)/Homer | 1e-6 | 0.0000 | 340.0 | 30.09% | 4735.7 | 23.32% |
| 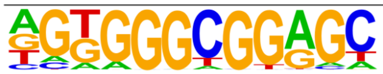 | Sp5(Zf)/mES-Sp5.Flag-ChIP-Seq(GSE72989)/Homer | 1e-5 | 0.0005 | 189.0 | 16.73% | 2462.9 | 12.13% |
| 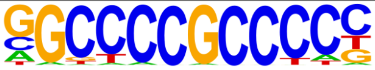 | Sp1(Zf)/Promoter/Homer | 1e-4 | 0.0016 | 68.0 | 6.02% | 708.4 | 3.49% |
| 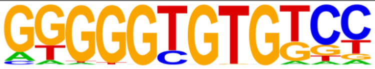 | KLF10(Zf)/HEK293-KLF10.GFP-ChIP-Seq(GSE58341)/Homer | 1e-3 | 0.0171 | 89.0 | 7.88% | 1082.6 | 5.33% |
| 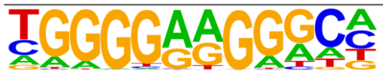 | ZNF467(Zf)/HEK293-ZNF467.GFP-ChIP-Seq(GSE58341)/Homer | 1e-3 | 0.0268 | 138.0 | 12.21% | 1861.5 | 9.17% |
| 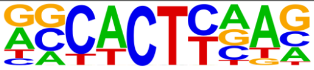 | Nkx2.1(Homeobox)/LungAC-Nkx2.1-ChIP-Seq(GSE43252)/Homer | 1e-2 | 0.0698 | 395.0 | 34.96% | 6233.4 | 30.70% |
| 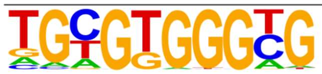 | Egr1(Zf)/K562-Egr1-ChIP-Seq(GSE32465)/Homer | 1e-2 | 0.0698 | 105.0 | 9.29% | 1395.4 | 6.87% |
| 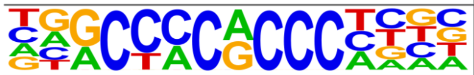 | KLF3(Zf)/MEF-Klf3-ChIP-Seq(GSE44748)/Homer | 1e-2 | 0.0698 | 108.0 | 9.56% | 1447.1 | 7.13% |
| 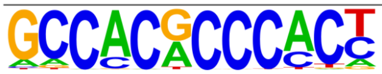 | Klf9(Zf)/GBM-Klf9-ChIP-Seq(GSE62211)/Homer | 1e-2 | 0.0698 | 76.0 | 6.73% | 959.9 | 4.73% |

**Figure S7. Sequences detected as enhancers in the MPRA are enriched for transcription factor binding site motifs compared to sequences without detected enhancer activity.** Analysis was performed using HOMER (*33*).

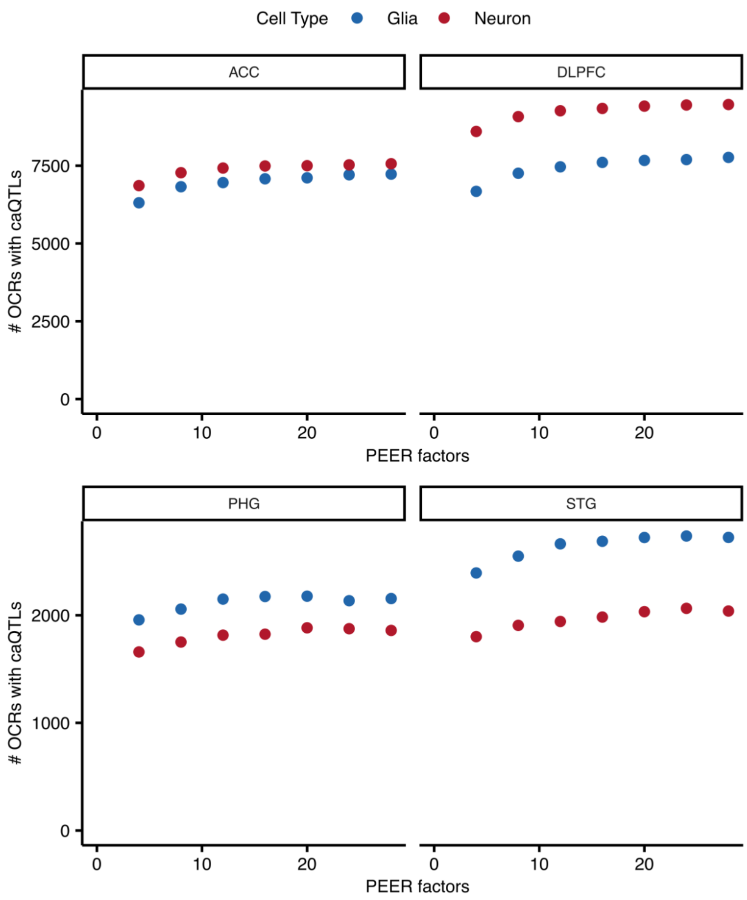


**Figure S8. PEER factors increase OCRs with detected caQTLs.**
caQTL analysis was performed using between 4 and 28 PEER factors, and the number of OCRs with genome-wide significant caQTLs was computed. The analysis was repeated for each cell type and brain region to choose the number of PEER factors for each subset of the data.

### Tables S1 to S2

**Table S1: Colocalization results for a) eQTL-caQTL-GWAS, b) caQTL-GWAS and c) eQTL_GWAS.**

Suppl_table_coloc.xlsx

**Table S2: SNPs with significant allelic effects by MPRA that are also significant by GWAS.** This number is small since the MPRA was designed to test eQTL rather than GWAS variants.

MPRA_GWAS_integration.xlsx

24. J. Nasser, D. T. Bergman, C. P. Fulco, P. Guckelberger, B. R. Doughty, T. A. Patwardhan, T. R. Jones, T. H. Nguyen, J. C. Ulirsch, F. Lekschas, K. Mualim, H. M. Natri, E. M. Weeks, G. Munson, M. Kane, H. Y. Kang, A. Cui, J. P. Ray, T. M. Eisenhaure, R. L. Collins, K. Dey, H. Pfister, A. L. Price, C. B. Epstein, A. Kundaje, R. J. Xavier, M. J. Daly, H. Huang, H. K. Finucane, N. Hacohen, E. S. Lander, J. M. Engreitz, Genome-wide enhancer maps link risk

variants to disease genes. *Nature*. **593**, 238–243 (2021).
